# Risk-based Decision Making in Rats: Modulation by Sex and Amphetamine

**DOI:** 10.1101/2020.03.09.981563

**Authors:** Dannia Islas-Preciado, Steven R. Wainwright, Julia Sniegocki, Stephane E. Lieblich, Shunya Yagi, Stan B. Floresco, Liisa A. M. Galea

**Affiliations:** Department of Psychology; Djavad Mowafaghian Centre for Brain Health; Graduate Program in Neuroscience. University of British Columbia, Vancouver, BC, V6T 1Z3, Canada

**Keywords:** amphetamine, decision-making, operant, gonadal hormones, 17β-estradiol, testosterone, cognition

## Abstract

Decision-making is a complex process essential to daily adaptation in many species. Risk is an inherent aspect of decision-making and it is influenced by gonadal hormones. Testosterone and 17β-estradiol may modulate decision making and impact the mesocorticolimbic dopamine pathway. Here, we explored sex differences, the effect of gonadal hormones and the dopamine agonist amphetamine on risk-based decision making. Intact or gonadectomised (GDX) male and female rats underwent to a probabilistic discounting task. High and low doses of testosterone propionate (1.0 or 0.2 mg) and 17β-estradiol benzoate (0.3 μg) were administered to assess acute effects on risk-based decision making. After 3-days of washout period, intact and GDX rats received high or low (0.5 or 0.125 mg/kg) doses of amphetamine and re-tested in the probabilistic discounting task. Under baseline conditions, males made more risky choices during probability discounting compared to female rats, particularly in the lower probability blocks, but GDX did not influence risky choice. The high, but not the low dose, of testosterone modestly reduced risky decision making in GDX male rats. Conversely, 17β-estradiol had no significant effect on risky choice regardless of GDX status in either sex. Lastly, a higher dose of amphetamine increased risky decision making in both intact males and females, but had no effect in GDX rats. These findings demonstrated sex differences in risk-based decision making, with males showing a stronger bias towards larger, uncertain rewards. GDX status influenced the effects of amphetamine, suggesting different dopaminergic regulation in risk-based choices among males and females.

## Introduction

We routinely face situations that require decisions entailing evaluation of varying costs and benefits associated with different actions. Among the different types of costs that may diminish the subjective value of larger or more preferred rewards, risk/reward decisions involve choices between options that have a lower risk but yield relatively inferior rewards, or more valuable rewards associated with some uncertainty as to whether they will be received (Wiehler and Peters, 2015). Dopamine plays an integral role in modulating cost/benefit decision making as pharmacological blockade of dopamine receptors reduced preference for larger rewards associated with various costs, including effort, delays and uncertainty (Denk et al., 2005; St. Onge et al., 2010; Nunes et al., 2010; Yohn et al., 2015).

The mesolimbic dopamine system is modulated by both sex hormones and sex (Becker, 2009; Morris et al., 2015; Yoest et al., 2014). Studies in humans have reported sex differences in different forms of risk-related decision making, which in turn may be driven by sex hormones (Op de Macks et al., 2016; Peper et al., 2013). Human males and females with high testosterone show more risk-related behaviour (Stanton et al., 2011) and high testosterone has been linked to poorer decision making in the Iowa gambling task (Reavis and Overman, 2001; van Honk et al., 2004). In addition, increased 17β-estradiol levels may be related to high self-reported risk-taking behaviour in females (Bröder and Hohmann, 2003; Sukolová and Sermány-Schuller, 2011). Thus, testosterone and 17β-estradiol modulate risk-taking behaviours in both human males and females (Op de Macks et al., 2016; Vermeersch et al., 2008a, b).

Preclinical studies in rats have also identified sex differences in risk preferences that may be related to sex hormones. For example, compared to females, male rats show a greater preference for larger rewards associated with a foot shock delivered in a probabilistic manner (Orsini et al., 2016). Likewise, treatment with supraphysiological doses of chronic testosterone in male rats further increases the preference for larger rewards associated with punishment (Cooper et al., 2014). On the other hand, using a probabilistic discounting task wherein rats chose between small/certain and larger/uncertain rewards (in the absence of any punishment), treatment with supraphysiological doses of chronic testosterone in adolescent male rats reduced risky choice (Wallin et al., 2015). This highlights that androgenic modulation of decision making can vary depending on the type of risk an animal faces (punishment or omission of the reward). However, it is not yet known whether 17β-estradiol or acute physiological doses of testosterone can also modulate risk-based decision making in males.

As alluded to above, sex differences in risk-related decision making may be mediated in part through hormonal modulation of dopamine activity. This may include alterations in dopaminergic tone and axon density within the nucleus accumbens and medial prefrontal cortex (Aubele and Kritzer, 2011; Dazzi et al., 2007; Saigusa et al., 1997), which play a key role in guiding risk/reward decision biases (St. Onge et al. 2010; St. Onge et al., 2012; Stopper et al., 2013; Jenni et al., 2017). 17β-Estradiol increases binding on D2 dopamine receptors and facilitates the excitability of dopaminergic transmission in females (Thompson and Moss, 1994; Becker and Hu, 2008). Testosterone increases dopamine synthesis (Purves-Tyson et al., 2012), the expression of D2 receptor mRNA in the midbrain, and reduced extracellular dopamine levels in the prefrontal cortex after gonadectomy in males (Aubele and Kritzer, 2011; Kritzer et al., 1999; Purves-Tyson et al., 2014). Thus, sex hormones modulate dopaminergic function in both males and females, which may in turn, influence risk/reward decision biases (Becker et al., 2012; Sotomayor-Zarate et al., 2014; Walker et al., 2017).

Although some studies have examined how exogenous testosterone may influence risk/reward decision making, considerably less is known about how reductions in testosterone may alter these functions. There is also a lack of data on how estrogens may influence these types of decisions and given that testosterone can be converted to 17β-estradiol, it is important to determine the effects of estrogens on decision making in males. In addition, increased preferences for risky rewards in males versus females have been reported when larger rewards are associated with punishment (Mitchell et al., 2011; Orsini et al., 2015; Simon et al., 2009), which may be related to sex differences in fear expression (Colon et al., 2018; Farrell et al., 2013; Gruene et al., 2015). Yet, how sex may influence decisions guided solely by reward uncertainty in the absence of potential punishment is unknown. To address these issues, we explore sex differences, the effects of gonadectomy, and the influence of testosterone or 17β-estradiol, on performance within a probabilistic discounting task, where rats chose between small/certain and larger/uncertain rewards. Additionally, we sought to determine whether gonadal hormone status may interact with manipulations of dopamine transmission in regulating risky choice, using acute treatment with amphetamine, which disrupts adjustments in choice in response to changes in reward probabilities (St. Onge et al., 2010). Here, we hypothesized that males would show greater preference for larger/risky rewards that would be heightened by testosterone and that gonadal hormone status would mediate the risky decisions made in male and female rats by enhancing the response to amphetamine in intact rats.

## Methods

### Animals

Twenty male and sixteen female Long Evans rats (Charles River Laboratories, Montreal, Canada) weighing 275-300 g and 200-250g, respectively, at the beginning of behavioural training were used for the experiment. On arrival, rats were given one week to acclimatize to the colony and food-restricted to 85-90% of their free-feeding weight for an additional one week before behavioural training. Rats were given ad libitum access to water for the duration of the experiment. Feeding occurred in the home cage at the end of the experimental day, and body mass was monitored daily to ensure a steady weight loss during food restriction and maintenance or weight gain for the rest of the experiment. All testing was in accordance with the Canadian Council of Animal Care and was approved by the Animal Care Committee of the University of British Columbia.

### Surgery

Gonadectomy (GDX) surgery was performed under isoflurane anesthesia (5% in oxygen during induction, 3% in oxygen during maintenance). Treatment groups were assigned according to their baseline choice performance over the last three days of training, such that there was no difference between groups in all blocks.

*Males* were either bilaterally castrated or received sham-castrations. For castrations, both testes were extracted through a small incision made at the posterior tip of the scrotum and were ligated with a monofilament suture. Sham operations involved incisions into the skin and muscle layers of the scrotum that were sutured without removing the testes. *Females* were either bilaterally ovariectomized or received sham-ovariectomy. For ovariectomy, both ovaries were extracted through a small horizontal incision in the abdominal wall directly over each ovary and were ligated with a monofilament suture. Sham operations involved incisions through the skin and abdominal muscle that were sutured without removing the ovaries. Prior to surgery, each rat was given a subcutaneous injection of Ketoprofen (5 mg/kg body mass) as an analgesic and immediately afterward, Flamazine cream (1% silver sulfadiazine) was applied to the incision. All rats received Ketoprofen 24 and 48 hours after surgery at 5 mg/kg subcutaneously (s.c.) as an analgesic. After surgery, rats were singly housed and one week was allowed for full recovery from surgery prior to further experimentation.

### Drug Treatments

Both GDX and sham *males* were assessed for changes in baseline choice performance following surgery. Upon returning to baseline performance, all rats received vehicle (sesame oil) or hormone treatment, as described previously on the experimental timeline. Between each hormone or drug treatment, baseline performance was also re-assessed. Single injections of testosterone propionate (0.2mg or 1mg, s.c.) and 17β-estradiol benzoate (0.3μg, s.c.) were used to assess their acute activational effects. Testosterone propionate was chosen as it has the fastest elimination half-life (0.8 days) in comparison to other esters (Behre et al., 2004) and produces steeper peaks (reviewed in Kornmann et al., 2009) to assess acute effects. Doses of hormones were chosen to mimic high and/or low physiological levels of these hormones (Chowdhury and Tcholakian, 1979; Holmes et al., 2002; Spritzer and Galea, 2007; Uban et al., 2012; Wainwright et al., 2016).

Both GDX and sham *females* were assessed for changes in baseline choice performance following surgery. Upon returning to baseline performance, all rats received hormone treatment accordingly to the experimental timeline. A single injection of 17β- estradiol benzoate (0.3μg) was used to assess their acute activational effects.

For both males and females, after completion of hormone treatments, we assessed the effects of amphetamine (0.125mg/kg or 0.5mg/kg i.p.) in both treatment groups (GDX and sham). I.P. administration of amphetamine was used because the dose-response and time-course of the effects of this drug on probabilistic discounting has been well-characterized (St. Onge and Floresco 2008; St. Onge et al., 2010). Each drug test consisted of a two-day sequence. The first day, rats were injected with saline, and the second day, with one of the two doses of amphetamine (counterbalanced). After each injection, rats were placed in their home cages and behavioural testing commenced 10 min later. After the first test sequence, rats were trained for three days before receiving another test sequence of saline and the other dose of amphetamine (based on St. Onge and Floresco, 2009). There were no differences in performance across the two saline injection tests, so the data from these tests were averaged in the analysis and compared to the two amphetamine challenges.

### Apparatus

Behavioural testing for all experiments was conducted in 12 operant chambers for the males and four operant chambers for the females (30.5 × 24 × 21 cm; Med-Associates, St Albans, VT) enclosed in sound-attenuating boxes in different rooms. The boxes were equipped with a fan to provide ventilation and to mask extraneous noise. Each chamber has two retractable levers, one located on each side of a central food receptacle where food reinforcement (45 mg; Bioserv, Frenchtown, NJ) was delivered via a pellet dispenser. The chambers were illuminated by a single 100-mA houselight located in the top center of the opposite wall to the levers. Locomotor activity was calculated by the number of photobeam breaks that occurred during a session as four infrared photobeams were mounted on the sides of each chamber. The chambers were connected to a computer through a digital interface that recorded all experimental data.

### Experimental timeline

The experimental timeline is shown in Figure 1. Each animal received Lever-Pressing Training 1 week after the initiation of food restriction. After 5-6 days of Lever-Pressing Training, animals were trained on the probabilistic discounting task to examine the baseline choice performance. After completion of the baseline training, half of the animals (equal sexes) were gonadectomised and half were given a sham surgery. Based on previous work, animals were given one week for recovery (Spritzer and Galea, 2007; Koss et al., 2018; Wagner et al., 2018) as this results in reduced hormone levels (Woolley and McEwen et., 1993; Kashiwagi et al., 2005) and then were retrained on the task to examine the baseline choice performance after the surgery. Two-weeks after gonadectomy, rats received vehicle treatment and then the next day, the first dose of hormone injections and the discounting test 3h, 24 and 48 h later to test rapid and delayed effects of the steroid hormones. Rats then were re-trained to baseline choice performance for 3-7 days to ensure baseline performance and then received a second dose of hormone and then a discounting test. The same procedure was repeated for the third dose of hormone injections. After completion of hormone treatment, rats received two doses of amphetamine and examined their choice performance. Vehicle treatments were always conducted 24h before hormone or drug treatments (Figure 1).

**Figure 1.**
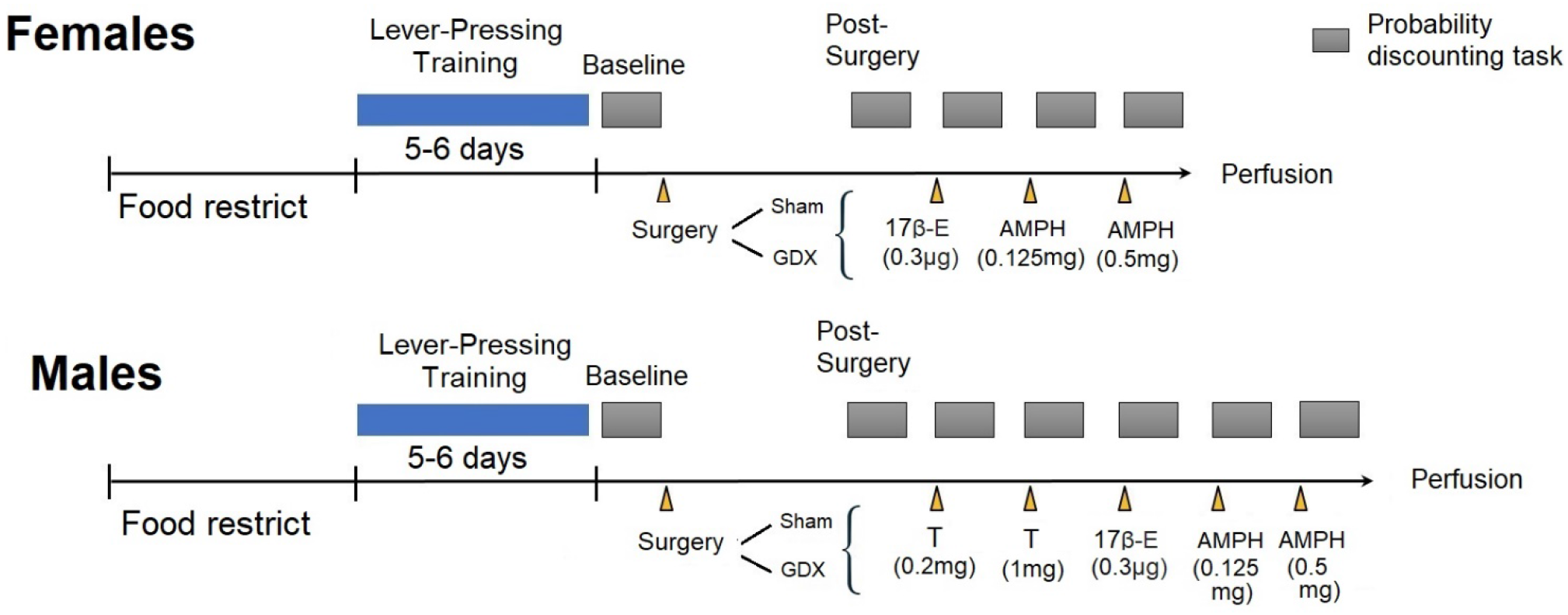
Experimental timeline. GDX=Gonadectomy, 17β-E= 17β- Estradio benzoate, Amph= Amphetamine, T= Testosterone

### Lever-Pressing Training

Our initial training protocols have been described previously (Stopper et al., 2013). Briefly, rats were first trained under a fixed-ratio schedule to a criterion of 60 presses in 30 min, first for one lever, and then repeated for the other lever (counterbalanced left/right between subjects). Rats were then trained on a simplified version of the full task - 90 trial sessions beginning with the levers retracted and the operant chamber in darkness, and each trial requiring them to press one of the two levers within 10s of its insertion for reinforcement of a single pellet delivered with 50% probability. If the rat did not respond to a lever within 10s after the lever was presented, the chamber was reset to the intertrial state until the next trial (omission). This procedure was used to familiarize the rats with the association of lever pressing and food delivery, as well as the probabilistic nature of the full task. In every pair of trials, the left or right lever was presented once, and the order within the pair of trials was randomized. Rats were trained for approximately 5-6 days to reach a criterion of less than ten omissions out of 90 trials.

### Probability Discounting Task

The task was modified from Cardinal and Howes’ (2005) procedure, which has been previously used to assess the role of dopaminergic and noradrenergic neurotransmission in risk-based decision making (St Onge and Floresco, 2009; Montes et al., 2015). Rats received daily sessions 6-7 days per week, consisting of 72 trials each day, separated into four blocks of 18 trials over 48min, where the probability of receiving a large reward decreased each block (100, 50, 25, 12.5%). A session began in darkness with both levers retracted (the intertrial state). A trial began every 40s with the insertion of one or both levers into the chamber 3s after the illumination of the house light. One lever was assigned as the Large/Risky lever, the other the Small/Certain lever. The combination of levers with the amount of reward given remained consistent throughout training but counterbalanced the position (left or right). If the rat did not respond by pressing a lever within 10s of lever presentation, both levers were retracted and the house light was turned off without any reward was given (omission). When a lever was chosen, both levers retracted. Pressing the Small/Certain lever always delivered one pellet with 100% probability, but pressing the Large/Risky lever delivered four pellets with 0.5 s apart but with a specific probability of the trial block. When food was delivered, the house light remained on for another 4s after a response was made, which was followed by reversion of the chamber to the intertrial state. The four blocks were comprised of 18 trials with eight forced-choice trials followed by ten free-choice trials. During a forced-choice trial, only one lever was presented (4 trials for each lever, randomized in pairs), allowing animals to learn the amount of food associated with each lever pressing and the respective probability of receiving reward over each block. During a free-choice trial, both levers were presented and the rat chose either the Small/Certain or the Large/Risky lever. The probability of receiving large reinforcement after pressing the Large/Risky lever was initially 100%, then 50%, 25%, and 12.5%, respectively, for each successive block. For each session and trial block, the probability of receiving the large reinforcement was drawn from a set probability distribution. Therefore, on any given day, the probabilities in each block may have varied slightly from the assigned probability, but on average across many training days, the actual probability experienced by the rat was approximate the set value. Using these probabilities, selection of the Large/Risky lever was more advantageous in the first two blocks (100% and 50% probability trial), and more disadvantageous in the last block (12.5%) compared to selection of the Small/Certain lever, whereas rats could obtain an equivalent number of food pellets after responding on either lever during the 25% block. Therefore, in the last three trial blocks (50%, 25% and 12.5%) of this task, selection of the larger reward option carried with it an inherent ‘‘risk’’ of not obtaining any reward on a given trial. Latencies to initiate a choice and overall locomotor activity (photobeam breaks) were also recorded. The criteria for the completion of training on the task was that rats chose the Large/Risky lever during the first trial block (100% probability) on at least 80% of successful trials.

### Data Analyses

The primary dependent measure of interest was the proportion of choices directed toward the large reward lever for each block of free-choice trials, factoring out trial omissions. For each block, this was calculated by dividing the number of choices of the large reward lever by the total number of successful trials (i.e., those where the rat made a choice). Data from each vehicle treatment test day were averaged. The effect of trial block was always significant (p<0.001) for the probabilistic discounting task and will not be discussed further. Repeated-measures analyses of variance were conducted with between-subject factors of sex (male, female) and surgery (sham, GDX) and within-subjects factors of block (12.5, 25, 50, 100% of probability of the large/risky reward), hormone dose (testosterone 0,0.2,1 mg or 17β- estradiol benzoate 0, 0.3 μg), time (pre-GDX, post-GDX) or amphetamine dose (0,0.125 0.5 mg/kg) were conducted using Statistica (v. 13, StatSoft, Tulsa, OK, USA). Post-hoc tests utilized Newman-Keuls comparisons, whereas a priori comparisons used a Bonferroni correction. Effect sizes are given as partial η_p_^2^ or Cohen’s *d* where appropriate. The significance level was set at α= 0.05. Response latencies and the number of trial omissions were analyzed with one-way repeated-measures ANOVAs.

Whenever a significant main effect of a treatment or group on probabilistic discounting was observed, we conducted a supplementary analysis to further clarify whether changes in choice biases were due to alterations in sensitivity to reward (win-stay performance) or negative-feedback (lose-shift performance) (Stopper et al., 2013). Animals’ choices during the task were analyzed according to the outcome of each preceding trial (reward or non-reward) and expressed as a ratio. The proportion of win-stay trials was calculated from the number of times a rat chose the large/risky lever after choosing the risky option on the preceding trial and obtaining the large reward (a win), divided by the total number of free-choice trials where the rat obtained the larger reward. Conversely, lose-shift performance was calculated from the number of times a rat shifted choice to the small/certain lever after choosing the risky option on the preceding trial and was not rewarded (a loss), divided by the total number of free-choice trials resulting in a loss. This analysis was conducted for all trials across the four blocks. We could not conduct a block X block analysis of these data because there were many instances where rats either did not select the large/risky lever or did not obtain the large reward at all during the latter blocks. These data were analyzed with multifactorial ANOVAs with Response type (win stay/lose shift) as one within-subject factor, drug/hormone treatment as another within-subject factor (when applicable) and Sex and Gonadectomy status as a between-subjects factor.

## Results

### Males showed greater bias for large/risky rewards versus females, that was not altered by gonadectomy

Figure 2 shows the proportion of risky choices made by male and female rats on the last three days of training pre-surgery (baseline, Fig. 2A) and for three days of training two weeks post-surgery (Fig. 2B). Analysis of the choice data at baseline revealed a sex X trial block interaction (F(3, 93)=3.5, p=0.018; η_p_^2^=0.10; Fig 2A). Males showed significantly greater choice of the large/risky lever in both the 25% (p<0.02, Cohen’s *d=* 0.563) and 12.5% (p<0.00018, Cohen’s *d=*0.645) probability trial blocks compared to females. There were no other significant main effects or interactions.

**Figure 2.**
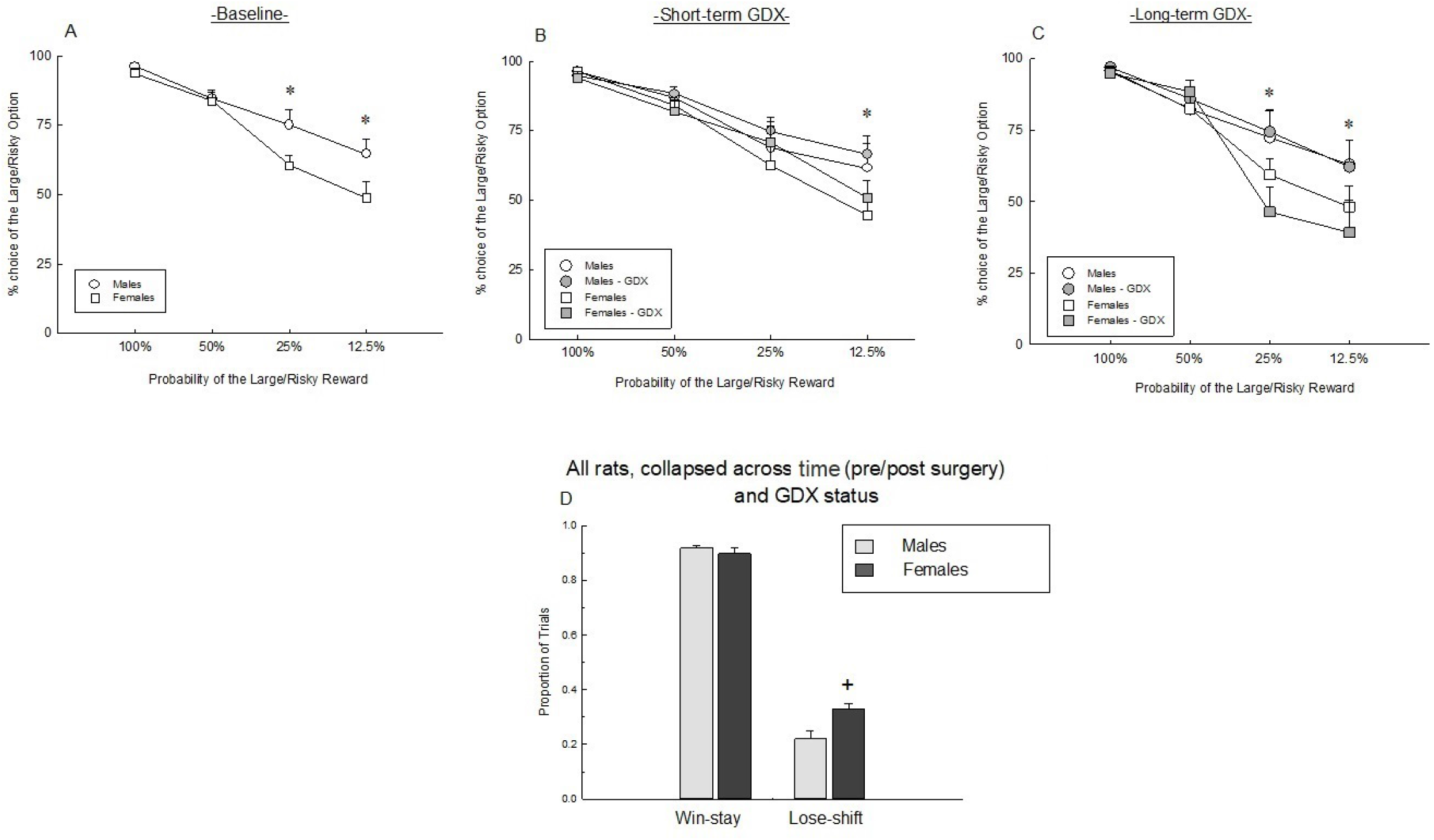
Group means (+ SEM) of the percentage choice of the Large/Risky lever during 100%, 50%, 25% and 12.5% trial blocks at baseline (2A) short-GDX (2B) and long-term GDX (2C) in male and female rats. Panel D shows the Win-stay and Lose-shift ratio during the probabilistic discounting task. n= 19 males, 16, females at baseline; n= 8 −10 per group in short and long-term GDX. * denotes p<0.05 vs females at 25% and 12.5% block regardless of GDX; + denotes p<0.05 vs males. Abbreviations: GDX, gonadectomy.

Gonadectomy (GDX, short-term – two weeks post-surgery) did not influence performance in this task compared to sham animals (all p’s> 0.55. Figure 2B). As previous studies have shown differences in performance between short versus long-term GDX (Bimonte-Nelson et al., 2003; Aubele and Kritzer, 2011), we conducted a further analysis to determine whether there were any surgery effects six weeks after GDX. However, even six weeks after GDX there were no significant effects of GDX relative to sham (all p’s >0.51. Figure 2 C), and males still made more choices on the risky lever in the 25% (Cohen’s *d*= 0.90) and 12.5% (Cohen’s d= 0.85) probabilities than females regardless of GDX (p’s <0.05) status; sex by block interaction: (F(3, 93)=5.463, p<0.0017, η_p_^2^=0.15).

The lower levels of risky choice observed in female rats under baseline conditions and post-GDX appeared to be associated with an increased tendency to shift to the small/certain lever after a non-rewarded risky choice (lose-shift behaviour). Although analysis of these data yielded a response type X sex interaction that only approached statistical significance (F(1,32)=3.36, p=0.076), an exploratory comparison confirmed that females displayed higher lose-shift ratios than males (F(1,34)=4.28, p=0.046 Fig 2D). There were no other interactions across pre/post-surgery phases or GDX status (all p’s>0.25). Thus, female rats were more sensitive to the ability of non-rewarded risky choices to influence subsequent choices away from the risky option compared to males.

At baseline females displayed a greater number of trial omissions (1.063 ± 0.3) than males (0.246 ± 0.1) main effect of sex F(1, 27)=4.929, p=0.035, η_p_^2^= 0.15) but there was no significant effect of GDX on the number of trial omissions (all p’s > 0.35). Females had significantly greater choice latency compared to males (main effect of sex F(1, 27)=4.886, p=0.036, η_p_^2^= 0.15), and there was a main effect of time (F(1, 27)=4.848, p=0.036, η_p_^2^= 0.15) showing choice latency to be significantly greater with time, following either GDX or sham surgery. As expected, females also showed significantly greater locomotor activity compared to males at baseline (main effect of sex F(1, 29)=20.409, p=0.00001, η_p_^2^= 0.41, and surgery had no significant effect on locomotor activity (all p’s > 0.11) (see Table 1).

**Table 1.** Effect of hormonal treatments on omissions, latency to decision-making and locomotor activity of sham and GDX males and females rats.

| *Group* | *Males Sham* |  |  | *Females Sham* |  |  |
| --- | --- | --- | --- | --- | --- | --- |
|  | *Omissions* | *Latency* | *Locomotor activity* | *Omissions* | *Latency* | *Locomotor activity* |
| *Baseline* | 0.2 ± 0.1 | 0.8 ± 0.1 | 1389.6 ± 83.8 | 1.1 ± 0.3<sup>b</sup> | 1.1 ± 0.1<sup>c</sup> | 2101.2 ± 174.1<sup>d</sup> |
| *Vehicle* | 0.1 ± 0.1 | 0.8 ± 0.1 | 1761.0 ± 220.3 | 2.3 ± 1 | 1.5 ± 0.2 | 1714.0 ± 223.6 |
| *T - 0.2 mg* | 0.5 ± 0.3 | 0.8 ± 0.2 | 1662.5 ± 180.4 | ----- | ----- | ----- |
| *T - 1 mg* | 0.2 ± 0.1 | 0.8 ± 0.1 | 1723.8 ± 179.5 | ----- | ----- | ----- |
| *17βE - 0.3 μg* | 0.8 ± 0.5 | 0.9 ± 0.2 | 1649.3 ± 136.7 | 3.7 ± 1.0 | 1.6 ± 0.2 | 1712.9 ± 244.0 |
| *Vehicle 24 hrs* | 0.6 ± 0.3 | 0.8 ± 0.2 | 1895.4 ± 148.9 | 4.4 ± 1.2 | 1.8 ± 0.2 | 1765.1 ± 217.4 |
| *T - 0.2 mg 24 hr* | 1.6 ± 0.9 | 1.0 ± 0.2 | 1516.6 ± 164.9<sup>a</sup> | ----- | ----- | ----- |
| *T - 1mg 24 hr* | 1.9 ± 0.9 | 1.0 ± 0.2 | 1509.8 ± 117.1<sup>a</sup> | ----- | ----- | ----- |
| *17βE - 0.3μg 24 hrs* | 0.8 ± 0.4 | 0.8 ± 0.2 | 1653.7 ± 147.5 | 2.9 ± 1.3 | 2.0 ± 0.2<sup>e</sup> | 1626.4 ± 160.1 |
|  | *Males GDX* |  |  | *Females GDX* |  |  |
|  | *Omissions* | *Latency* | *Locomotor activity* | *Omissions* | *Latency* | *Locomotor activity* |
| *Short tem* | 0.1 ± 0.1 | 0.9 ± 0.1 | 1263.7 ± 122.5 | 1.0 ± 0.6 | 1.2 ± 0.2 | 1562.3 ± 101.4 |
| *Long term* | 0.8 ± 0.6 | 0.8 ± 0.1 | 1651.7 ± 171.6 | 4.2 ± 2.0 | 1.8 ± 0.3 | 1486.6 ± 102.3 |
| *Vehicle* | 0.3 ± 0.2 | 0.8 ± 0.1 | 1244.3 ± 80.2 | 2.0 ± 1.0 | 1.2 ± 0.2 | 1439.1 ± 192.6 |
| *T - 0.2 mg* | 0.2 ± 0.2 | 0.8 ± 0.1 | 1211.7 ± 107.4<sup>aa</sup> | ----- | ----- | ----- |
| *T - 1 mg* | 0.0 ± 0.0 | 0.8 ± 0.2 | 1206.9 ± 158.3<sup>aa</sup> | ----- | ----- | ----- |
| *17βE - 0.3 μg* | 1.1 ± 1.1 | 1.0 ± 0.2 | 1059.9 ± 132.2 | 1.8 ± 0.8 | 1.3 ± 0.2 | 1423.5 ± 169.9 |
| *Vehicle 24 hrs* | 0.3 ± 0.2 | 0.8 ± 0.1 | 1309.5 ± 210.7 | 2.3 ± 1.0 | 1.4 ± 0.2 | 1443.0 ± 154.9 |
| *T - 0.2 mg 24 hr* | 0.0 ± 0.0 | 0.8 ± 0.1 | 1158.9 ± 131.7<sup>aa</sup> | ----- | ----- | ----- |
| *T - 1 mg 24 hr* | 0.0 ± 0.0 | 0.8 ± 0.1 | 1010.9 ± 126.9<sup>aa</sup> | ----- | ----- | ----- |
| *17βE - 0.3 μg 24 hrs* | 0.8 ± 0.7 | 1.0 ± 0.2 | 990.3 ± 135.4 | 1.5 ± 0.6 | 1.5 ± 0.2<sup>e</sup> | 1513.0 ± 136.3 |

<sup>a</sup> denotes p<0.05 testosterone – 24h vs vehicle (reduced locomotor activity after 24h of testosterone treatment)
<sup>aa</sup> denotes p<0.01 GDX vs sham (GDX males had reduced locomotor activity than sham males treated with testosterone)
<sup>b</sup>denotes p<0.05 females vs males (females had grater omissions than males)
<sup>c</sup> denotes p<0.05 females vs males (females had greater latency than males)
<sup>d</sup> denotes p<0.001 females vs males (females had grater locomotor activity than males)
<sup>e</sup> denotes p<0.05 17β-estradiol – 24h vs vehicle (increased latency after 24h of 17β-estradiol treatment)

### High, but not low, Testosterone treatment modestly reduced risky decision making in the GDX males only in the 12.5% probability block

Figure 3 shows the acute effects of low (0.2 mg/kg) and high (1.0 mg/kg) dose of testosterone on risk-based decisions in sham (3A) and GDX (3B) male rats. The high, but not low, dose of testosterone reduced proportion of risky choices in the lowest probability block (12.5%; p<0.002, Cohen’s *d=*0.815) compared within the GDX males but not in the sham males (p=0.40; dose X block X surgery F(6, 90)=2.025, p=0.07, η_p_^2^=0.12). The high dose of testosterone increased the probability of a lever press for the 25% block (p<0.04), but this is no longer significant after a Bonferroni correction. There was a main effect of blocks as expected (F(6, 45)=22.119, p<0.0001, η_p_^2^=0.596) but no other significant main or interaction effects (p’s> 0.11).

**Figure 3.**
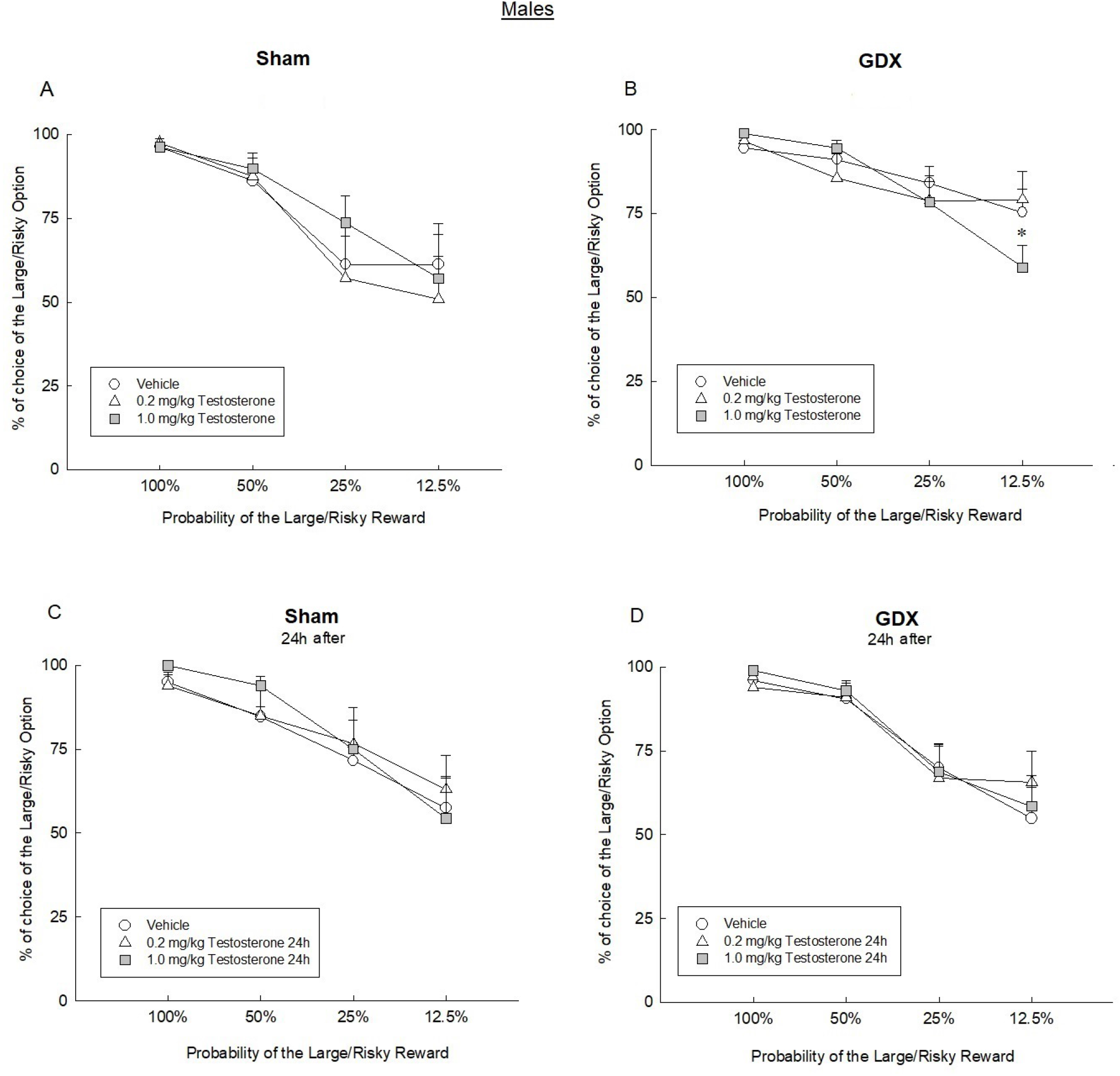
Group means (+ SEM) of the percentage choice of the Large/Risky lever during 100%, 50%, 25% and 12.5% trial blocks in both, males sham (2A) and GDX (2B) treated with low and high dose of testosterone, and 24 hours after testosterone administration in sham (2C) and GDX (2D) rats. n= 8-10 per group. * denotes p<0.05 vs vehicle. Abbreviations: GDX, gonadectomy

There was no significant effect of testosterone treatment (at low or high dose) on the number of trial omissions or choice latency (all p’s > 0.07; Table 1). Figure 3C and D shows the risky choices at 24 h after treatment with low and high dose of testosterone. As it can be seen, no significant effect on the risk-based choices was observed after testosterone administration (main effect of dose: p=0.571, main effect of block F(3, 54)=32.802, p<0.0001), all other main or interaction effects p’s>0.22). However, both the low (Cohen’s *d*= 0.45) and high (Cohen’s *d*= 0.73) doses of testosterone treatment decreased locomotor activity after 24h of testosterone treatment (p <0.035; Table 1). There was also a main effect of surgery, where GDX males had significantly reduced locomotor activity than sham males (F(1, 14)=9.246, p<0.009, η_p_^2^= 0.40) (see Table 1).

### 17β-estradiol treatment did not influence risky decision making in males or females

Figure 4 shows the effect of 17β-estradiol benzoate on the risky choices across different groups. Figure 4A and 4B partition the data based on GDX status collapsed across sexes and panels C-F partition these data further by sex. There were no significant effects of 17β-estradiol treatment on risky decision making in either sex (no significant main or interaction effects with sex (all p’s >0.17). There was a significant interaction of 17β-estradiol dose X surgery X block effect (F(6, 180)=2.412, p=0.029, η_p_^2^=0.074), but post-hoc tests failed to find anything meaningful. There was a significant main effect of block (F(3, 90)=70.774, p<0.0001, η_p_^2^=0.702) but no other main or interaction effects (all p’s >0.17). After 24 hrs of 17β-estradiol treatment, no changes were observed in the risky choices neither in male nor female rats independently of GDX status. There was also no significant effect of 17β-estradiol treatment on the number of omissions in either males or females (all p’s >0.39) (Table 1). However, 17β-estradiol treatment significantly increased choice latency in females 24h after treatment (F(1, 29)=7.829, p=0.009, η_p_^2^=0.21) (table 1). There were no other significant effects on choice latency (all p’s >0.15) (Table 1).

**Figure 4.**
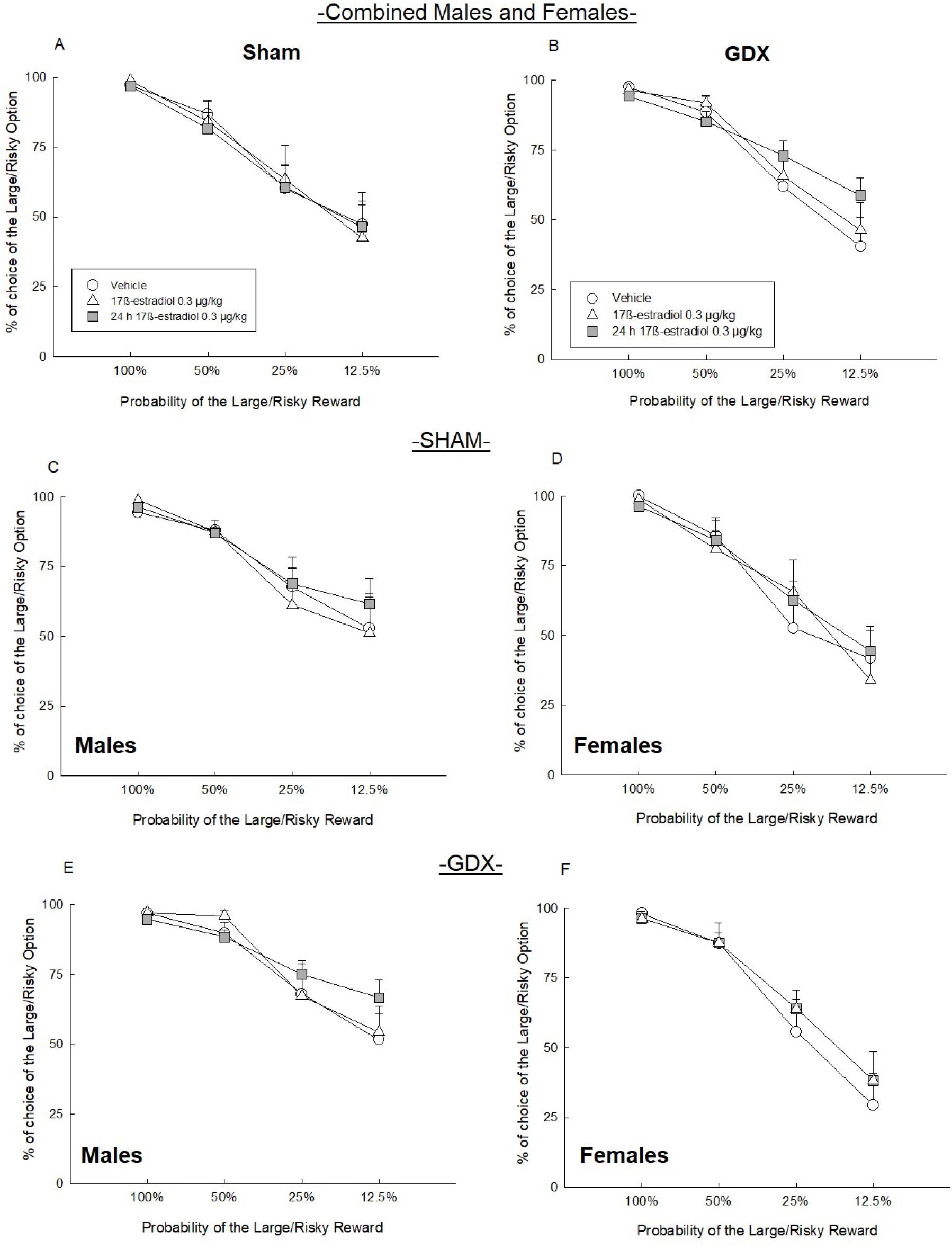
Group means (+ SEM) of the percentage choice of the Large/Risky lever during 100%, 50%, 25% and 12.5% trial blocks in combined males and females sham (5A) and GDX (5B) rats, males sham only (5C) and female sham only (5D), treated with 17β-estradiol and 24 hours after 17β-estradiol treatment. Abbreviations: GDX, gonadectomy.

### Amphetamine increased risky choice in both males and females, but this effect was blunted by GDX

Figure 5 shows the effect of amphetamine (AMPH) on risky choices across the different groups and sexes. One female sham rat displayed a marked increase in omissions over all trial blocks, and its choice data were excluded from the analysis. Analysis of these data yielded a significant AMPH dose X block interaction (F (6,186)=5.805, p<0.001) and a dose X block X GDX interaction (F (6,186)=3.15, p<0.01), but no other 3 or 4 way-interactions with the Sex Factor (all p’s>0.10). Figure 5A and B partition the data based on GDX status, collapsed across sexes and panels C-F partition these data further by sex. Simple main effects revealed that in sham animals, the 0.5 mg/kg dose increased risky choice in the 25% and 12.5% probability blocks (p<0.05), whereas the 0.125 mg/kg dose only increased risky choice in the 12.5% block (p<0.05). In GDX rats, this effect was markedly blunted, wherein the 0.5 mg/kg dose of amphetamine did not induce a statistically significant increase in risky choice during any block (all p’s>0.15). AMPH treatment also did not disrupt the inherent sex difference (block X sex interaction (F (3,93)= 8.71, p<0.0001, η_p_^2^=0.22). These findings reveal that the effects of AMPH on risk-based decision making is comparable in both males and females, and that indicates that GDX in both sexes attenuates the ability of amphetamine to alter risky choice.

**Figure 5.**
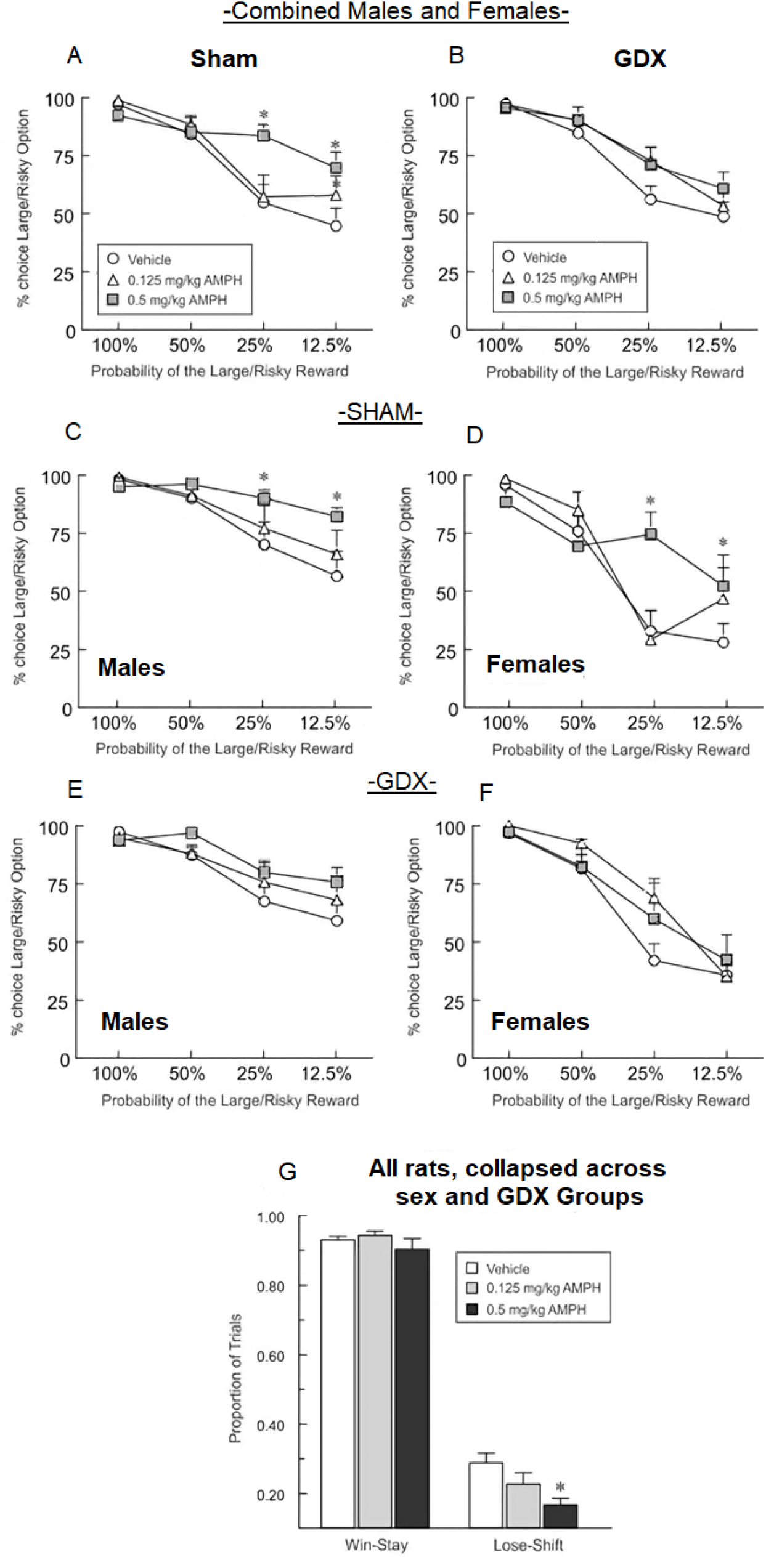
Group means (+ SEM) of the percentage choice of the Large/Risky lever during 100%, 50%, 25% and 12.5% trial blocks in combined males and females sham (5A) and GDX (5B) rats, males sham only (5C), females sham only (5D), male GDX only (5E), female GDX only (5F) treated with a low (0.125 mg/kg,) or high (0.5 mg/kg) dose of amphetamine. Panel E shows the Win-stay and Lose-shift ratio during the probabilistic discounting task. n= 8-10 per group. * denotes p<0.05 vs vehicle. Abbreviations: GDX, gonadectomy, AMPH, amphetamine.

The increase in risky choice induced by AMPH across all groups was driven by a reduction in lose-shift tendencies. Analysis of these data revealed a significant dose X response type interaction (F(2,26)= 3.57, p=0.034) but no other interactions by dose with the sex or GDX factors (all p’s>0.10). Simple main effects analysis further confirmed that AMPH treatments did not alter win-stay behaviour (p>0.30), but the 0.5 mg/kg dose significantly reduced lose-shift behaviour (p<0.001; Fig 5G). There was also a significant response type X sex interaction (F(1,31)=4.66, p=0.048) indicating that across all drug doses and conditions, females displayed greater lose-shift tendencies (0.27 +/−0.03) vs males (0.19+/−0.02) but no differences in win-stay behaviour (females=0.91+/−0.02; males 0.94 +/−0.01).

With respect to other performance measures, the 0.5 mg/kg dose of AMPH increased the number of omissions (dose X sex X surgery interaction F(3, 93)=8.851, p=0.006, η_p_^2^=0.22), wherein sham females had a greater number of omissions than all other treatment groups despite the interaction (all p’s< 0.0002). In contrast, the 0.125 mg/kg dose of AMPH decreased the number of omissions (main effect of treatment F(1, 29)=5.520, p=0.026, η_p_^2^=0.16). The low dose of AMPH decreased choice latency in females (treatment by sex interaction F(1, 30)=8.900, p=0.006, η_p_^2^=0.23) but females still showed greater choice latency than males (all p’s< 0.02). The high dose of AMPH increased choice latency in females (main effect of sex (F(1, 28)=20.429, p=0.0001, η_p_^2^=0.42) and the high dose of AMPH tended to induce a longer latency in sham females in comparison to female GDX (sex by surgery interaction (F(1, 28)=3.797, p=0.06, η_p_^2^=0.12). A low dose of AMPH produced no significant effect on locomotor activity, although there was a trend (F(1, 23)=3.728, p=0.066, η_p_^2^=0.14). As expected, high dose AMPH increased locomotor activity in all treatment groups (day by sex interaction (F(1, 25)=16.770, p=0.0004, η_p_^2^=0.40), and increased locomotor activity in females more than males (p < 0.005) (Table 2).

**Table 2.** Effect of low and high dose of amphetamine on omissions, latency to decision-making and locomotor activity of sham and GDX males and females rats

| Group | Males Sham |  |  | Females Sham |  |  |
| --- | --- | --- | --- | --- | --- | --- |
|  | Omissions | Latency | Locomotor activity | Omissions | Latency | Locomotor activity |
| Vehicle 0.125 mg/kg | 0.1 ± 0.1 | 0.8 ± 0.2 | 1128.5 ± 117.0 | 5.3 ± 3.0 | 2.6 ± 0.8 | 1405.3 ± 276.6 |
| AMPH 0.125 mg/kg | 0.0 ± 0.0<sup>a</sup> | 0.8 ± 0.2 | 1622.3 ± 189.3 | 6.0 ± 5.5 | 1.8 ± 0.6<sup>e</sup> | 1372.0 ± 152.5 |
| Vehicle 0.5 mg/kg | 0.2 ± 0.2 | 0.7 ± 0.2 | 1764.8 ± 136.1 | 1.7 ± 0.3 | 2.0 ± 0.2 | 1531.7 ± 140.9 |
| AMPH 0.5 mg/kg | 0.0 ± 0.0<sup>a</sup> | 0.8 ± 0.2 | 2184.7 ± 278.6<sup>b</sup> | 8.0 ± 5.0<sup>d</sup> | 2.2 ± 1.0<sup>f</sup> | 2345.3 ± 377.7<sup>b,g</sup> |
|  | Males GDX |  |  | Female GDX |  |  |
| Vehicle 0.125 mg/kg | 0.8 ± 0.7 | 0.8 ± 0.1 | 1651.7 ± 171.6 | 4.2 ± 2.0 | 1.7 ± 0.3 | 1486.6 ± 102.3 |
| AMPH 0.125 mg/kg | 0.6 ± 0.3<sup>c</sup> | 0.8 ± 0.1 | 1039.9 ± 92.3 | 1.0 ± 0.7<sup>c</sup> | 1.5 ± 0.3<sup>e</sup> | 1322.2 ± 193.5 |
| Vehicle 0.5 mg/kg | 0.6 ± 0.6 | 0.8 ± 0.2 | 1138.4 ± 100.9 | 3.0 ± 1.6 | 1.6 ± 0.3 | 1386.3 ± 217.9 |
| AMPH 0.5 mg/kg | 1.2 ± 0.7 | 0.9 ± 0.1 | 1804.5 ± 288.7<sup>b</sup> | 1.5 ± 1.3 | 1.5 ± 0.3<sup>f</sup> | 2376.5 ± 237.9<sup>b,g</sup> |

<sup>a</sup> denotes p<0.05 AMPH 0.125 and 0.5 mg/kg vs vehicle (reduced omissions after AMPH treatment)
<sup>b</sup> denotes p<0.0005 AMPH 0.5 mg/kg vs vehicle (increased locomotor activity by AMPH)
<sup>c</sup> denotes p<0.05 AMPH 0.125 mg/kg vs vehicle (reduced omissions after AMPH treatment)
<sup>d</sup> denotes p<0.05 AMPH 0.5 mg/kg vs vehicle, (increased omissions after AMPH treatment)
<sup>e</sup> denotes p<0.05 AMPH 0.125 mg/kg vs vehicle (reduced latency after AMPH treatment)
<sup>f</sup> denotes p<0.0005 females vs males, (greater latency in females than in males after AMPH 0.5 mg/kg)
<sup>g</sup> denotes p<0.005 females vs males (greater locomotor activity in females than in males treated with AMPH 0.5 mg/kg)

## Discussion

The present study revealed notable sex differences during risk/reward decision making, in that male rats made more risky choices on a probabilistic discounting task compared to females. However, these sex differences were not driven by adult levels of gonadal hormones, as gonadectomy did not alter risky choice in either sex, nor did they alter sex differences in risk/reward decision making. Acute treatment with a higher dose of testosterone propionate (3 h) modestly reduced risky choice in GDX, but not intact, male rats. Furthermore, a low physiological dose of 17β-estradiol benzoate had no significant effect on decision making of either males and females regardless of their gonadectomy status. Lastly, amphetamine increased risky decision making of both male and female rats in a dose-dependent manner, and this effect of amphetamine was more pronounced in rats with intact gonads compared to GDX animals.

### Male rats made more risky choices during the probabilistic discounting task than female and gonadectomy did not influence their risk-related decision making

Male rats made more risky choices when the probability of obtaining the food reward was relatively low (25% or 12.5%) compared to female rats. Importantly, there were no sex differences in choice of the large/risky reward when the probability of obtaining a larger food reward was 50 or 100%, suggesting that the lower levers of risky choice observed in females was unlikely due to reduced preference for larger rewards. These results are consistent with a previous study using another type of risky decision-making task in which the risk of larger reward was associated with a negative consequence of footshocks (Orsini et al., 2016). On the other hand, our results are slightly inconsistent with Georgiou et al. (2018) who used a rat version of the Iowa Gambling Task (IGT). They found that male rats chose the most advantageous choice more often than females. Whereas in the present study, we found that females showed lower levels of risky choice in the lower probability of high reward trial blocks, suggesting that female rats avoided risky choices. This sex difference appeared to be driven in part by an increased tendency by females to shift their choice after encountering an occasional loss regardless of whether it was in the long-term advantageous or disadvantageous, compared to males, suggesting that females may be more loss averse. This tendency was also observed in human studies evaluating risk/loss aversion using the Risky-Gains Task (Lee et al., 2009) and the Cambridge Gambling Task (van den Bos et al., 2014).

Sex differences have also been observed among different types of decision-making in rodent models. Adult female mice displayed less impulsive behaviour than males by reducing their preference for a larger, delay reward in intertemporal decision-making (Koot et al., 2009; Perry et al., 2007). Along similar lines, sex differences have been proposed in effortful in decision-making, as females may exert more effortful control than males (Bjorklund and Kipp, 1996), but may also show a reduced preference for larger rewards associated with a greater effort cost (Uban et al., 2012). Our results here add evidence of sex differences in decision making that encompass risky choices, and that males made more risk-based choices in comparison to females when animals are choosing between certain and uncertain rewards without encountering potential punishment.

Interestingly, a recent study did not observe sex differences in rats performing a probabilistic discounting task similar to the one used here (Braunscheidel et al., 2019), which contrasts with our findings. These differential effects may be related to several procedural differences between the two studies, including shipping stress (greater impacts on gonadal hormone responses if shipping during puberty compared to shipping during adulthood: Laroche et al., 2009), strain of rats used (Long Evans versus Sprague Dawley), housing conditions, and the type of reinforcer used (solid reward pellets versus sucrose solution). Furthermore, the study by Braunscheidel and colleagues (2019) used a task comprised of more trials 5 blocks, including an additional 6.25% probability block which produced greater discounting compared to the performance of rats in our study. These seemingly minor differences may have occluded an ability to detect differences in discounting between males and females. Nevertheless, the results of these two studies show that detection of sex differences in risky choice may be dependent on the particular experimental procedures used.

Gonadectomy influences some aspects in decision-making in males (Tan and Vyas, 2016; Wallin-Miller et al., 2017) and females (Uban et al., 2012). However, in the present study, gonadectomy did not influence risky choices in either males or females. This discrepancy can be due to different types of costs that were associated with larger rewards across different tasks (Tan and Vyas, 2016; Uban et al., 2012), or if the response was elicited by another stimulus (Wallin-Miller et al., 2017), as in the latter study, risky choices were elicited by ethanol and this could impact the mesolimbic reward pathway to alter risk-based decisions. The idea that sex-hormone regulation of decision making can differ depending on the type of cost associated with larger/preferred rewards is further bolstered by the findings of studies examining how stress affects these processes. Specifically, acute stress can influence certain forms of cost/benefit decision making without affecting others, in that acute stress can reduce preference for larger rewards associated with a greater effort cost, but not those that are delayed or delivered in a probabilistic manner (Shafiei et al., 2012; Bryce and Floresco, 2016; Bryce et al., 2020).

Intriguingly, there are examples of sex differences in the effects of acute stress on learning as acute stress facilitates associate learning in males, but impairs it in females (Wood and Shors, 1998). The present data further highlight that the hormonal and neurochemical modulation of different forms of cost/benefit decision making can vary based on the type of cost being evaluated. Moreover, gonadal hormones may influence food intake and impact on the behavioural performance in a food-based reward task. However, our results do not indicate that gonadectomy altered trial omissions, which are an index of motivation to perform the task.

### Testosterone and estradiol effects on risk/reward decision making-interactions with gonadectomy

We found that an acute high dose of testosterone induced a minimal effect by reducing risky choices only during the 12.5% probability trial block compared to vehicle-treated GDX rats. This is partially consistent with a previous study using a similar probability discounting task showing that chronic supraphysiological testosterone (7.5 mg) throughout adolescence (5-14 weeks old) reduced risky choice compared to vehicle-treated intact rats during 50%, 25% and 0% probability trial blocks (Wallin et al., 2015). The larger effect of testosterone in that study may be due dose, age of testosterone administration and/or gonadal status. Intriguingly, studies with supraphysiological doses of testosterone using the punishment (foot shocks) discounting task (Cooper et al., 2014), effort discounting (Wallin et al., 2015) or delay discounting task (Wood et al., 2013) showed that these large doses of chronic testosterone increased choice of large reward option. These data further highlights that testosterone can differentially affect decisions involving a choice between larger and smaller rewards in a manner that is critically dependent on the type of cost associated with these rewards. Indeed, studies show that testosterone can inhibit stress effects, including on memory, in males (Viau and Meaney, 1996; Panizzon et al., 2018). Thus, testosterone reduces the impact of punishments, delays or effort costs on diminishing the subjective value of larger rewards, resulting in an increased preference for these rewards. Conversely, increased testosterone activity appears to augment sensitivity to reward uncertainty, reducing choice of larger/uncertain rewards.

Androgen receptors (ARs) are located throughout the mesolimbic regions and on dopaminergic neurons in the prefrontal cortex (PFC) and dorsolateral striatum (Tobiansky et al., 2018), as these areas are reported as critical brain regions involved during risk/reward decision making (Orsini et al., 2015; Worthy et al., 2016). Indeed, systemic testosterone, but not 17β- estradiol, decreased the basal dopamine levels in the PFC in male rats (Aubele and Kritzer, 2011) and burst firing in the VTA was androgen, but not 17β-estradiol, sensitive (Locklear et al., 2017). This is important as testosterone can be converted to 17β-estradiol via aromatase and thus can modulate dopaminergic systems via its actions on AR or estrogen receptors (ER). Furthermore, systemic administration of an aromatase inhibitor (letrozole) decreased dopaminergic turnover in the PFC in both male and female rats (Kokras et al., 2018). Thus, androgens exert an important role in attenuating the disruptions in cortical innervations of dopamine induced by gonadectomy in male rats (Kitzer, 2000). Our observation that treatment with testosterone, but not 17β- estradiol, altered risk/reward decision-making in males adds additional support to the notion that that testosterone may alter these processes via AR but not ER and may do so by modulating mesocorticolimbic DA activity system. However, our results should be interpreted with caution as we only utilized one dose of 17β-estradiol, and it is possible with other higher doses we may see different effects in males.

We found no significant effect of ovariectomy or 17β-estradiol on risk-based decision making in female rats. These findings are consistent with evidence that estrous cycle did not influence risk-based decision making where risky choice is coupled with footshock (Orsini et al., 2016) or in the IGT (Georgiu et al., 2018). Conversely, a previous report showed that OVX rats made more effort for a lager reward, an effect that is not observed in intact animals and is reversed by 17β-estradiol (Uban et al., 2012), suggesting that 17β-estradiol may influence some aspects of decision making involving effort costs, but not those involving reward uncertainty. However, as noted it is possible higher doses may modify risky decision-making.

Interestingly, previous studies have found that 17β-estradiol (5 μg or 10 μg) can potentiate the release of dopamine in the dorsal striatum with amphetamine (Song et al., 2019), and in the nucleus accumbens (NAc) with cocaine (Tobiansky et al., 2016). Yet, in our case 17β- estradiol treatment did not alter risky choice, which can also be altered by pharmacological increases in dopamine transmission (St. Onge et al., 2009). Alternatively, 17β-estradiol may be able to augment drug-induced dopamine release and related behaviours, but on its own, might be insufficient to alter behaviourally evoked changes in dopamine (St. Onge et al., 2012). On the other hand, the fact that we observed that amphetamine increased risky choice in both sexes but only in intact and not in GDX rats, suggests that gonadal hormones can influence the ability of amphetamine to modulate risk/reward decision making. Collectively these studies suggest that although a low dose of 17β-estradiol may not directly influence risky choice, it may interact with the effects of dopaminergic drugs to alter reward-related decision biases. Supporting this, 17β- estradiol enhanced the amphetamine-induced place preference (Silverman and Koenig, 2007) and OVX decreased quinpirole specific binding in the NAc, which is reversed by 17β-estradiol administration (Le Saux et al., 2006).

### Amphetamine enhanced risky decision-making in both male and female rats, and this effect was stronger in sham rats compared to gonadectomised rats

Consistent with previous reports (St. Onge and Floresco, 2009; Floresco and Whelan, 2009; St. Onge et al., 2010), amphetamine treatments increased risky choice, and this effect was comparable across male and female rats. This general effect was associated with a reduction in negative feedback sensitivity, as rats were less likely to shift choices after a non-rewarded risky choice. This amphetamine-induced increase of risky decision during the probability discounting task is likely driven by activation of D1 and D2 receptors in different terminal regions (St. Onge and Floresco, 2009; St. Onge et al., 2011; Stopper et al., 2013; Pes et al., 2017). However, the ability of amphetamine to alter risky choices was not observed in GDX rats relative to those with intact gonads. The enhancing effect of gonadal hormones on amphetamine-induced risky decision-making is likely due to modulation of different dopaminergic systems between the two sexes. Indeed, cocaine-induced increase of dopamine levels in the dorsolateral striatum was enhanced by 17β-estradiol in female rats but not in males (Cummings et al., 2014). On the other hand, castration has been reported to enhance amphetamine-mediated dopamine release (Hernandez et al., 1994). Our experimental design did not detect statistical interactions between sex and gonadectomy status on the effects of amphetamine. Thus, future experiments will gain relevance to elucidate how gonadal hormones may exert influence on the ability of stimulant drugs to alter risk/reward decision making between sexes. It is also notable that, in addition to dopamine, amphetamine may alter other monoaminergic systems (Dawson et al., 2003; dela Pena et al., 2015) as well as glutamatergic transmission (Wolf et al., 2000). Moreover, glutamate pharmacological manipulations influence the preference for larger rewards in effort and delay discounting tasks (Floresco et al., 2008). Therefore, other systems may be playing a role in the interaction between amphetamine and gonadal hormones in males and females. This being said, the ability of AMPH to increase risky choice in males was completely abolished by treatment with dopamine receptor antagonists (St. Onge and Floresco, 2009), suggesting that this changes in DA release may be of primary importance in the effects of AMPH reported here.

## Conclusion

Sex differences exist in the cognitive manifestations of disorders involving the mesolimibic dopamine pathway such as schizophrenia, impulsive disorders and Parkinson’s disease, with males showing greater cognitive disruption. In the present study, we found that females were less likely to choose the riskier options than males, which was associated with increased sensitivity to loss. Surprisingly, gonadectomy did not influence risky decision-making in either sex under baseline conditions but did alter the manner in which amphetamine can influence action selection. These results suggest that dopaminergic regulation of risk-related behaviour is different depending on background hormone levels.

## Acknowledgments

We gratefully acknowledge support from the Natural Sciences and Engineering Research Council of Canada (NSERC) grant to LAMG (2018-04301) and SBF (2018-04295). DIP was funded by Consejo Nacional de Ciencia y Tecnología (Mexico) and SY was funded by a Killam Doctoral Award from the University of British Columbia.

